# A spatial transmission model for evaluating interventions during highly pathogenic avian influenza outbreaks

**DOI:** 10.64898/2026.10.14.758777

**Authors:** Fanny Bergström, Edward M. Hill, Cecilia Rago, Christopher N. Davis, Michael J. Tildesley, Robin N. Thompson

## Abstract

During infectious disease outbreaks, epidemiological models can be used to project future outbreak dynamics and to explore scenarios under different control measures. Here, we present the spatially explicit avian influenza transmission model that we developed during the 2026 WiLiMan-ID consortium Highly Pathogenic Avian Influenza Modelling Challenge, which involved evaluating control measures (preventive and reactive culling, confinement and restocking bans) during a hypothetical avian influenza outbreak. We show that the control measures that were expected to reduce transmission effectively varied between different outbreak stages. For example, policy-makers could choose whether to focus culling efforts on: i) faster reactive culling and no preventive culling; or ii) slower reactive culling and preventive culling. Relatively early in the outbreak, when the number of available culling teams was limited, our results indicate that the two culling strategies were projected to lead to similar numbers of premises with notified infections over the following 28 days. Later in the outbreak, with more culling teams available, the second strategy was projected to lead to substantially fewer notified premises than the first strategy. In addition, we demonstrate that whether a strategy is beneficial or detrimental depends on policy-makers’ objectives. For example, preventive culling may reduce transmission, but at the cost of culling poultry on premises without infections, illustrating a trade-off between limiting transmission and the cost of interventions. Our results highlight the importance of choosing policy objectives carefully and reassessing interventions at different outbreak stages to ensure that control policies remain appropriate for the current epidemiological situation.

## 1 Introduction

Highly pathogenic avian influenza (HPAI) represents a major threat to both wild birds and the poultry industry. In addition, infections have occurred in mammals, including dairy cattle in the USA (Rawson et al., 2025; Pekar et al., 2026). Since 2020, H5 lineage viruses have been responsible for widespread outbreaks affecting large numbers of commercial poultry premises, resulting in substantial economic losses and poultry mortality (Harvey et al., 2023; World Organisation for Animal Health, 2026). Control of HPAI in poultry typically relies on stringent interventions affecting the poultry industry, which can include culling, movement restrictions, confinement, restocking bans and vaccination (Vergne et al., 2024). Choosing which interventions to prioritise is challenging because the effectiveness of different measures depends on a wide range of factors, including the speed and spatial scale of transmission, the spatial distribution of poultry premises and the operational resources available for disease control.

Mathematical modelling is an essential tool for informing infectious disease outbreak control strategies (Heesterbeek et al., 2015; Thompson, 2020). Models that are fitted to outbreak data can be used not only to generate forwards projections of disease incidence during an outbreak based on current transmission patterns, but also to generate alternative projections with different control interventions in place (Kaminsky et al., 2019; Dankwa et al., 2022; Loo et al., 2024). In the context of livestock disease, the 2001 foot-and-mouth disease outbreak in the UK exemplifies the use of epidemiological models to guide control policy (Keeling et al., 2001; Ferguson et al., 2001; Kao, 2002; Woolhouse, 2003), and epidemiological models have been applied subsequently in the context of HPAI (Walker et al., 2012; Hill et al., 2017, 2018; Lambert et al., 2023; Davis et al., 2026). For example, previous mathematical modelling work has demonstrated the importance of proactive surveillance for controlling HPAI in poultry in Bangladesh (Hill et al., 2018) and shown that protective measures on poultry premises (e.g., enhanced biosecurity or vaccination) can reduce the numbers of premises with infections during HPAI outbreaks in the UK (Davis et al., 2026). Other recent research includes analyses of the transmission dynamics of the first HPAI outbreak in a commercial poultry farm in Brazil (Cardenas et al., 2026), the impact of outdoor farming, farm size and farm density on HPAI outbreaks in the Netherlands (Chanchaidechachai et al., 2026) and the spatial risk of HPAI in France (Artois et al., 2025).

Although models can be used to compare and rank possible control strategies to support policy decision making, the effectiveness of specific interventions is unlikely to remain constant over the course of an outbreak. Epidemiological conditions change as an outbreak progresses. For example, the availability of susceptible hosts to infect may vary (this could be due to susceptible depletion or because the location of the outbreak shifts to an area with different host density (Tildesley et al., 2010)) or operational constraints associated with implementing interventions may become more or less restrictive. Consequently, interventions that are effective early in an outbreak may become less effective later, while alternative strategies may become more favourable. In addition, the effectiveness of an intervention depends on the policy objective used to evaluate its success; interventions that perform well for one objective will not necessarily perform well for another (Probert et al., 2016; Thompson et al., 2026). To improve the effectiveness of outbreak control, it is crucial to understand how the relative performance of interventions varies over time according to the specified policy objective.

Here, we investigate these issues in the context of HPAI transmission during the hypothetical outbreak studied as part of the WiLiMan-ID consortium HPAI Modelling Challenge (Hayes et al., 2026a). During the challenge, modelling teams from around the world independently developed models to test different control measures and formulate policy recommendations. As the Oxford-Warwick-Liverpool (OWL) team, we developed a spatially explicit stochastic HPAI transmission model, building on previous research by some of the team members (Retkute et al., 2018; Hill et al., 2017, 2018; Davis et al., 2026). We used the model to evaluate interventions at different stages of the outbreak. In this article, we show that the model-projected performance of specific interventions changed during the outbreak and depended on the policy objective under consideration. For epidemiological models to effectively support decision making, our findings underscore the importance of continually assessing control strategies throughout outbreaks and explicitly defining policy objectives.

## 2 Methods

### 2.1 HPAI Challenge dataset

We analysed the synthetic outbreak dataset provided by the HPAI Modelling Challenge organisers. The outbreak took place on a fictional island (Jolly Island) with area 33,667 km^2^. Jolly Island has 9,160 poultry premises, split into five production types: layer chickens (441 premises), broiler stage one chickens (1,248 premises), broiler stage two chickens (2,746 premises), organic reared ducks (1,034 premises) and conventionally reared ducks (3,691 premises). A two-stage broiler production system divides the chicken growing cycle into an initial brooding phase (stage 1, typically days 1 to 21) and a final growing-finishing phase (stage 2, from day 22 to market weight).

During the outbreak, infections were detected on 560 premises in total in the period from 22 December 2025 to 11 April 2026. The outbreak was split into three phases (Fig. 1A). In Phase 1, the daily number of newly notified premises grew exponentially. In Phase 2, it continued to rise until late January 2026 and then declined. In Phase 3, it continued to decline.

**Figure 1:**
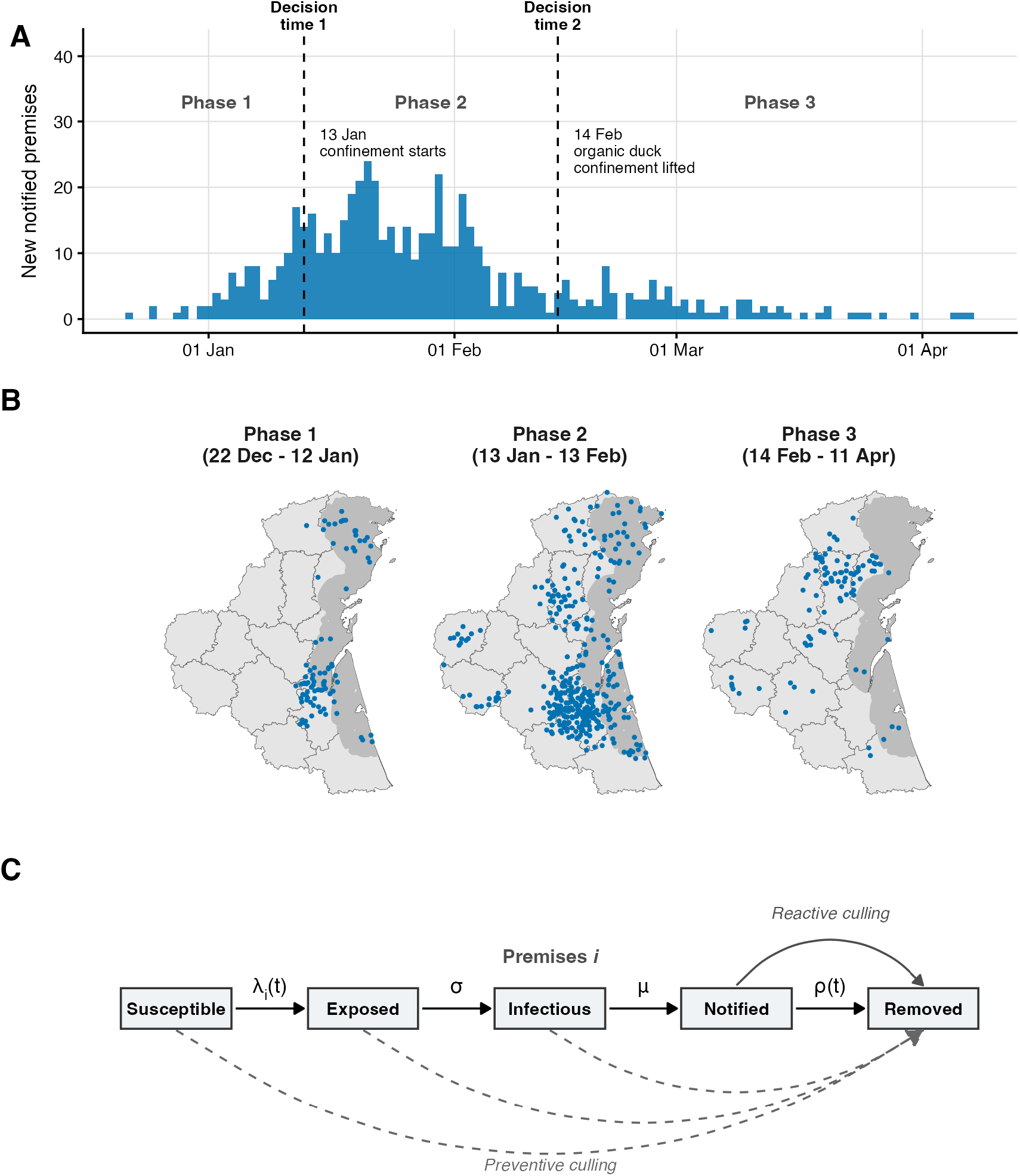
The hypothetical HPAI outbreak on Jolly Island and the compartmental HPAI transmission model structure. **(A)** Daily counts of newly notified premises on Jolly Island over the course of the outbreak (22 December 2025 to 11 April 2026), which is split into three outbreak phases. The two decision times (times at which we projected the effects of different control measures over a 28-day period) are also indicated by the dashed vertical lines; these decision times mark the boundaries between the phases. **(B)** Locations of premises with notified infections in each of the three outbreak phases (blue points); the dark grey shaded region indicates the high-risk zone in which imported infections were more likely. **(C)** The compartmental structure of our transmission model, in which the host unit is an individual poultry premises. Dashed arrows indicate that a premises may become removed due to preventive culling, bypassing the intermediate infection stages.

The control measures in place varied between the three outbreak phases. During Phase 1, reactive culling (culling of all poultry on premises with notified infections) was implemented, as was preventive culling of premises within one km of any premises with detected infections. Preventive culling was initiated when infections were confirmed on the premises that triggered the preventive cull. Throughout the outbreak, regulated zones were declared around each premises with detected infections for 28 days, comprising a 3 km protection zone and a 10 km surveillance zone (distinct from the high-risk zone for wild-bird incursions; see below); within these zones, poultry movements were restricted with premises only permitted to move poultry for slaughter. Culling was constrained by the limited availability of culling teams and there were no confinement measures in place in Phase 1. During Phase 2, reactive culling continued, while preventive culling was targeted solely at premises with chickens within three km of any premises with detected infections (regardless of the species on that premises), reflecting the assumption by policy-makers that premises with chickens were more likely to experience infections than premises with ducks. Additional culling teams were deployed progressively from the beginning of Phase 2, allowing more rapid implementation of culling, although the scale of this increase was only reported to modelling teams after Phase 2 (see Section 2.3). Confinement was also introduced on all premises with broiler stage two chickens or organic ducks at the start of Phase 2. In Phase 3, the same control measures remained in place as in Phase 2, except that confinement continued only for broiler stage two chickens.

Data were provided describing the locations and production types of each premises on Jolly Island, as well as the number of poultry on each premises (in periods in which the relevant premises was active). In addition, there were data describing the movement of poultry between premises and periods of inactivity (when no poultry were present on a given premises). For premises with detected infections, the following key dates relating to the outbreak response were available: the date on which infections were suspected on the premises, the date on which infections were confirmed and the date on which poultry on the premises were culled (as a result of either preventive or reactive culling).

### 2.2 Transmission model

Extending the model presented by Davis et al. (2026), we used a spatially explicit, stochastic, discrete-time HPAI transmission model. At any time, each premises was in one of five compartments: susceptible (*S*; no infected poultry on the premises), exposed (*E*; some poultry on the premises are infected, but not yet infectious so there is no risk of transmission to other premises), infectious (*I*; infectious poultry are on the premises), notified (*N* ; infections have been detected on the premises and HPAI has been confirmed, but culling has not yet taken place) or removed (*R*; all poultry on the premises have been culled). We used this model to represent transmission in all three outbreak phases, but we assumed some model parameters took different values in each phase to reflect transmission and the interventions that were in place at different stages of the outbreak.

In our model, if premises *i* is susceptible (i.e., there are no infections on the premises), the infectious pressure exerted on that premises on day *t* is

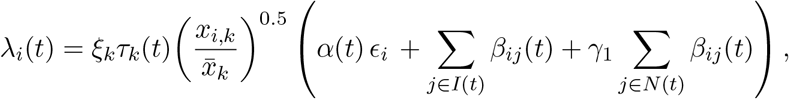

in which

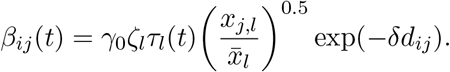

The three summed terms within the brackets in the expression for *λ_i_*(*t*) represent infectious pressure from three distinct sources. The first term, *α*(*t*)*ɛ_i_*, represents infection incursions into premises from the environment; for example, due to infected wild birds arriving on Jolly Island.

The second and third terms represent infectious or notified premises (*j*), respectively, increasing the risk of infections occurring on premises *i* (this could be due, for example, to lateral spread between premises or to vectored transmission between premises by wild birds). The subscripts *k* and *l* in the equations above refer to the production types of the susceptible premises (*i*) and infectious or notified premises (*j*), respectively.

The value of *ɛ_i_* depends on whether premises *i* lies within the high-risk zone (shown as the dark grey shaded area in Fig. 1B; the location of this zone was provided by the challenge organisers and reflects locations in which importations of infection from outside Jolly Island are most likely). We set *ɛ_i_* = *ɛ*_hrz_ if the premises is in the high-risk zone and *ɛ_i_* = *ɛ*_non-hrz_ if it is not. The variable *α*(*t*) reflects temporal variations in the risk of incursions of infection into Jolly Island during the transmission season: this parameter took the value one before 1 February 2026, and then declined exponentially to 0.008 on 28 February 2026 (and was fixed at zero thereafter). This schedule follows the information provided to modelling teams at the end of Phase 1. Specifically, modelling teams were told that, each year, most migratory wild birds arrive on Jolly Island in early winter, but that limited migration into the island continues through to the end of February.

The parameter *ξ_k_* represents the relative susceptibility of premises of production type *k* (where this parameter takes the value one for broiler stage one chickens). We set *τ_k_*(*t*) according to whether production type *k* was subject to confinement at time *t* (it was assumed to take the value 0 *< τ <* 1 if confinement was in place, and one otherwise). The parameters *x_i,k_* and *x̄_k_* represent the number of birds on premises *i* and the mean size of premises with production type *k*, respectively, so that the susceptibility of premises *i* increases with the number of birds on the premises. The exponent of 0.5 in the (*x_i,k_/x̄ _k_*)^0.5^ term ensures that the increase in susceptibility with premises size is sublinear; for example, a premises with twice as many birds of the same type as another premises is more susceptible, but not twice as susceptible.

The parameter *γ*_0_ determines the extent of between-premises transmission (again, either directly or vectored), and *γ*_1_ determines the relative infectious pressure generated on other premises by notified premises (compared to infectious premises). The dependence of this infectious pressure on the size of the infectious or notified premises acts in a similar fashion to the dependence of susceptibility on premises size. Confinement also affects the infectious pressure exerted by infectious or notified premises. The value of *ζ_l_* represents the relative infectious pressure generated by premises of production type *l* (this parameter also takes the value one for broiler stage one chickens). Finally, the parameter *δ* sets the rate at which the infectious pressure decays with the distance *d_ij_* between infectious or notified premises *j* and susceptible premises *i*.

A premises *i* that is susceptible at the beginning of day *t* becomes infected (and enters the *E* compartment of the model) on that day with probability 1 *−* exp(*−λ_i_*(*t*)). Exposed premises spend a mean period of *σ* days in the *E* compartment before becoming infectious, and infectious premises spend a mean period of *µ* days in the *I* compartment before notification, with the durations spent in each of these compartments sampled from independent probability distributions (see Table S1 for details; we also list all model parameters in this table).

Notified premises are culled after a mean delay of *ρ*(*t*) days, with *ρ*(*t*) taking the value *ρ*_1_ during Phase 1 (before 13 January 2026) and *ρ*_2_ from the start of Phase 2 onwards (from 13 January 2026). In addition to reactive culling of notified premises, preventive culling could be undertaken on a premises before the premises enters the *N* compartment, in which case the premises transitions to the *R* compartment after a mean delay of *ρ*(*t*) days. In addition to the delay to culling, the culling capacity (the number of premises that could be culled on any single day; *C*(*t*)) was known. This capacity was 15 premises per day before 13 January 2026, increased linearly to 100 premises per day between 13 and 30 January 2026, and remained at 100 premises per day thereafter (Table S1). When generating projections, if culling was due to occur on more than *C*(*t*) premises on a particular day, then culling was only implemented on *C*(*t*) premises, with the remaining premises instead being culled (and transitioning to the *R* compartment) on the next possible day. We assumed that reactive culling is generally prioritised over preventive culling, so that preventive culling occurred on any given day only after reactive culling had been undertaken.

All model parameters that are listed as “Fitted” in Table S1 were estimated by fitting the model to the outbreak data in a Bayesian framework using Stan (Carpenter et al., 2017). For each premises with detected infections, the unobserved day of infection (the transition from *S* to *E*) and the unobserved day of infectiousness onset (the transition from *E* to *I*) were marginalised out of the likelihood by summing over possible combinations of these days, with each term in the sum weighted by the probability of that combination (according to the latent and infectious period distributions). In the likelihood, the infectious pressure exerted by each premises with detected infections was calculated over a fixed window. This window began on the date on which infection was suspected (or seven days before the notification date for premises with no recorded suspicion date) and ended when culling started. We estimated the parameter values of the model using Markov chain Monte Carlo (MCMC; specifically using Stan’s No-U-Turn sampler), running four chains with 1,000 warm-up iterations and 1,000 sampling iterations per chain. We assessed convergence using the Gelman-Rubin statistic and the effective sample size (see Appendix B).

### 2.3 Counterfactual simulations and intervention comparison

We ran model simulations under a range of interventions to compare the projected effectiveness of different control strategies. Simulations were generated starting from either the beginning of Phase 2 (decision time 1; see Fig. 1A) or the beginning of Phase 3 (decision time 2), and were run for a period of 28 days. In each simulation, we sampled parameter values from the joint posterior distribution. We ran 1,000 simulations for each phase and intervention combination.

To compare individual simulations under different interventions, we simulated outbreaks using the Sellke algorithm (Sellke, 1983). For each simulation, we sampled a set of per-premises infection thresholds, *Q_i_ ∼* Exp(1), together with each premises’ latent and infectious periods and its baseline notification-to-culling period (these premises-specific periods were used if the relevant premises became infected). We reused these same values under every strategy compared for that specific simulation (although, as described below, the notification-to-culling period was reduced by one day in scenarios with faster reactive culling). Under the Sellke algorithm, a susceptible premises becomes infected at the first time *t* at which its accumulated force of infection (over time) exceeds its infection threshold (i.e., *_u≤t_ λ_i_*(*u*) *≥ Q_i_*), so that an intervention that lowers *λ_i_* either delays the time at which premises *i* becomes infected or prevents premises *i* becoming infected entirely (if the infection threshold is never exceeded). For a single simulation, differences in outcomes under different interventions thereby reflect only the effects of the interventions, as opposed to being due to stochasticity (Andersson and Britton, 2000).

In each simulation window, we ran simulations under a baseline scenario and under a range of counterfactual scenarios. The baseline scenario reflects the control policy communicated to modelling teams at the decision time from which the simulation was run. Hence, the baseline simulations represent the expected interventions as they were understood at the decision time. In the Phase 2 window, this differs from the policy that was actually implemented in two respects. First, the culling capacity in the Phase 2 window simulations was *C*(*t*) = 15 premises per day for all 28 days. At the first decision time, modelling teams were only informed that deployment of additional culling teams was being considered; the scale and timing of the increase described in Section 2.2 were only reported at the second decision time. Second, the Phase 2 baseline simulations do not include confinement, since the implementation of confinement was not described to modelling teams until the second decision time; confinement therefore enters the Phase 2 simulations only as a counterfactual intervention. In contrast to Phase 2, the interventions in the Phase 3 baseline simulations match those that were implemented in that phase. To check that the model can reproduce the observed outbreak dynamics, we also ran projections in which the culling capacity, confinement and preventive culling were set according to the interventions that were actually implemented, and compared these simulations with the observed daily notifications in each simulation window (Appendix C).

The interventions evaluated via counterfactual simulations comprised four different types: reactive culling, preventive culling, confinement and restocking bans (Table 1). For scenarios involving faster reactive culling, we reduced the notification-to-culling period for every premises by one day. For scenarios involving restocking bans, we used data provided by the challenge organisers describing periods when each premises was active or inactive. Whenever a premises was due to become active following a period of inactivity, we prevented this if a restocking ban was in place and at least one premises within a specified radius (either three km or ten km) had been notified (and entered the *N* compartment of the model) since the start of the simulation. Premises on which culling was undertaken during a simulation were not restocked for the rest of that simulation.

**Table 1:** Interventions evaluated in counterfactual simulations of the transmission model.

| Intervention | Description |
| --- | --- |
| Reactive culling | Culling of all poultry on premises with notified infections. |
| Preventive culling | Culling of all poultry on premises within a specified radius of any premises with notified infections. |
| Confinement | Housing restriction (requiring poultry to be kept indoors), thereby reducing the susceptibility and infectious pressure corresponding to all premises of the confined poultry types. |
| Restocking ban | Ban on repopulating premises (within a specified radius of any notified premises). |

### 2.4 Objectives of interventions

A range of metrics can be used to quantify the impact of an outbreak and to compare the effectiveness of different control strategies. One natural metric is the total number of notified premises, which provides a measure of the disease-induced impact of the outbreak. By comparing this quantity between simulations conducted under different control strategies, the extent to which each strategy reduces transmission can be assessed.

An alternative metric is the total number of affected premises, defined as premises that either have notified infections or on which preventive culling occurs. This metric captures both the direct impact of infection and the burden associated with preventive culling (whereby poultry must be culled even if infection has not been detected on the premises). A strategy that reduces the number of notified premises may not necessarily reduce the total number of affected premises. In most of our analyses, we compare control strategies using these two metrics. However, we also investigate how the preferred strategy changes when different relative weights are assigned to the burden of control interventions compared to the disease burden. The aim of this is not to provide a precise economic valuation of an outbreak or its control measures, but rather to illustrate how the policy objective (specifically, the relative preference for reducing disease or reducing the burden of interventions) can change which strategy may be preferred by policy advisors.

We consider two such comparisons. In our Phase 2 simulations, we vary the relative weight assigned to confinement and define the overall outbreak “cost” as the sum of the number of notified premises and the weighted number of premises subject to confinement. The weighted number of premises subject to confinement is the product of the number of premises subject to confinement and the per-premises confinement cost. Similarly, in our Phase 3 simulations, we instead vary the relative weight assigned to preventive culling and define the outbreak cost as the sum of the number of notified premises and the weighted number of premises on which preventive culling is undertaken. We use these analyses to show that different assumed control costs can lead to different optimal control strategies (i.e., different control strategies corresponding to the lowest overall outbreak cost).

## 3 Results

### 3.1 Intervention effectiveness differs between outbreak phases

We used MCMC to estimate the transmission model parameter values from the outbreak data. Posterior distributions are shown in Fig. S1 and the median parameter value estimates are given in Table S3. We then ran simulations of the fitted model to investigate the projected effects of different counterfactual control measures at different stages of the outbreak (specifically, in 28-day periods at the start of Phases 2 and 3 as described above).

In our simulations at the start of Phase 2, replacing preventive culling with faster reactive culling (by one day) was only projected to change the number of notified premises slightly: the median change relative to the baseline policy was *−*0.7% with faster reactive culling rather than preventive culling (with a 95% prediction interval (PI) of (*−*3.5%, 1.2%); Fig. 2A, Table 2).

**Figure 2:**
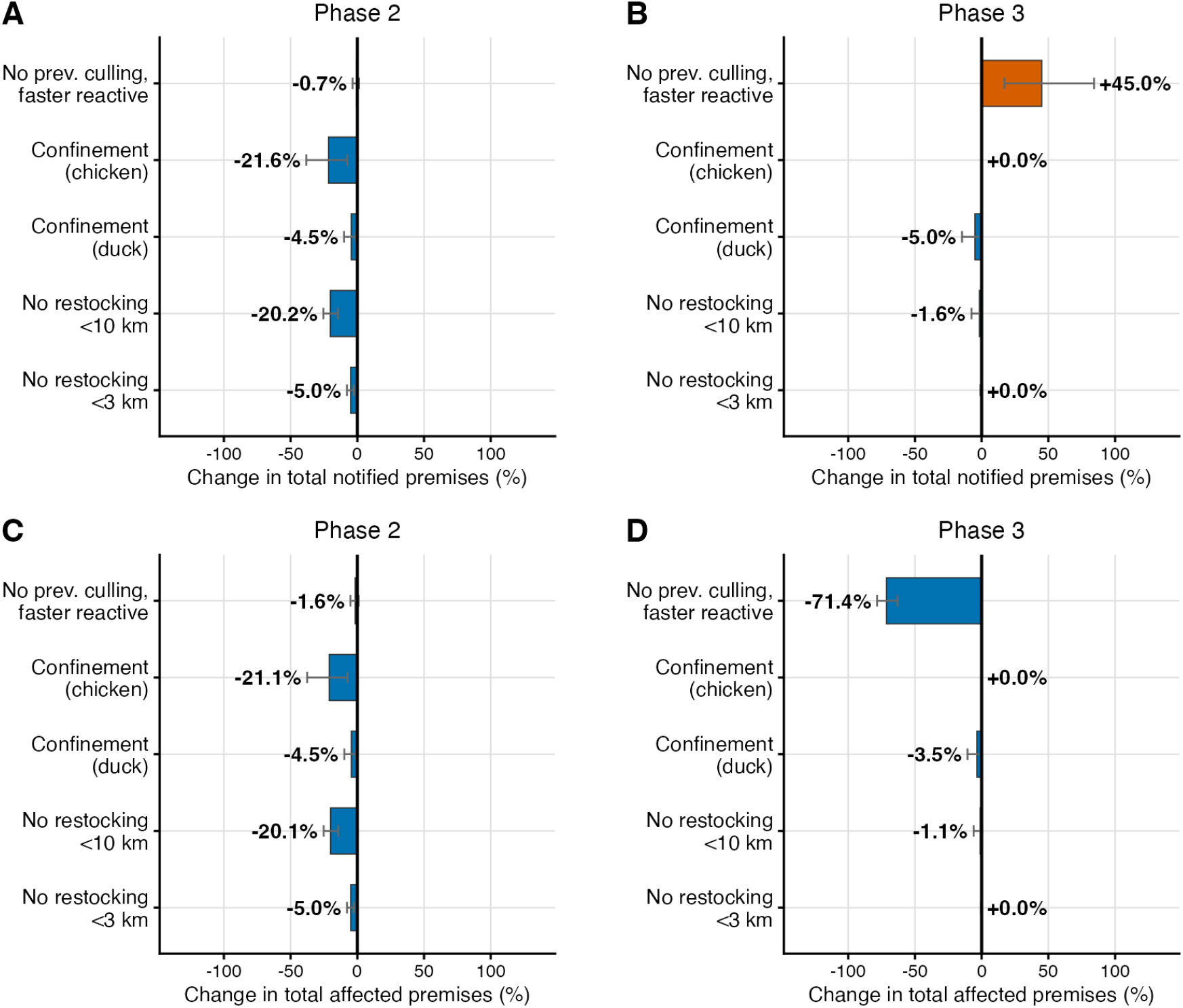
Projected epidemiological impacts of potential control policy changes relative to the baseline policy at each decision time. (A) Change in the total number of notified premises in the Phase 2 simulation period under different control interventions (relative to the baseline intervention scenario, which reflects the control policy communicated to modelling teams at decision time one). (B) Change in the total number of notified premises in the Phase 3 simulation period under different control interventions (relative to the baseline intervention scenario, which reflects the control policy communicated to modelling teams at decision time two). (C) Change in the total number of affected premises (defined as premises with notified infections or that were subject to preventive culling) in the Phase 2 simulation period under different control interventions (relative to the baseline intervention scenario, which reflects the control policy communicated to modelling teams at decision time one). (D) Change in the total number of affected premises (defined as premises with notified infections or that were subject to preventive culling) in the Phase 3 simulation period under different control interventions (relative to the baseline intervention scenario, which reflects the control policy communicated to modelling teams at decision time two). In all panels, each bar shows the median percentage change and error bars indicate the central 95% PI. Bar colour indicates whether the median change was positive (orange) or negative (blue). Simulations were run for 28 days.

**Table 2:**
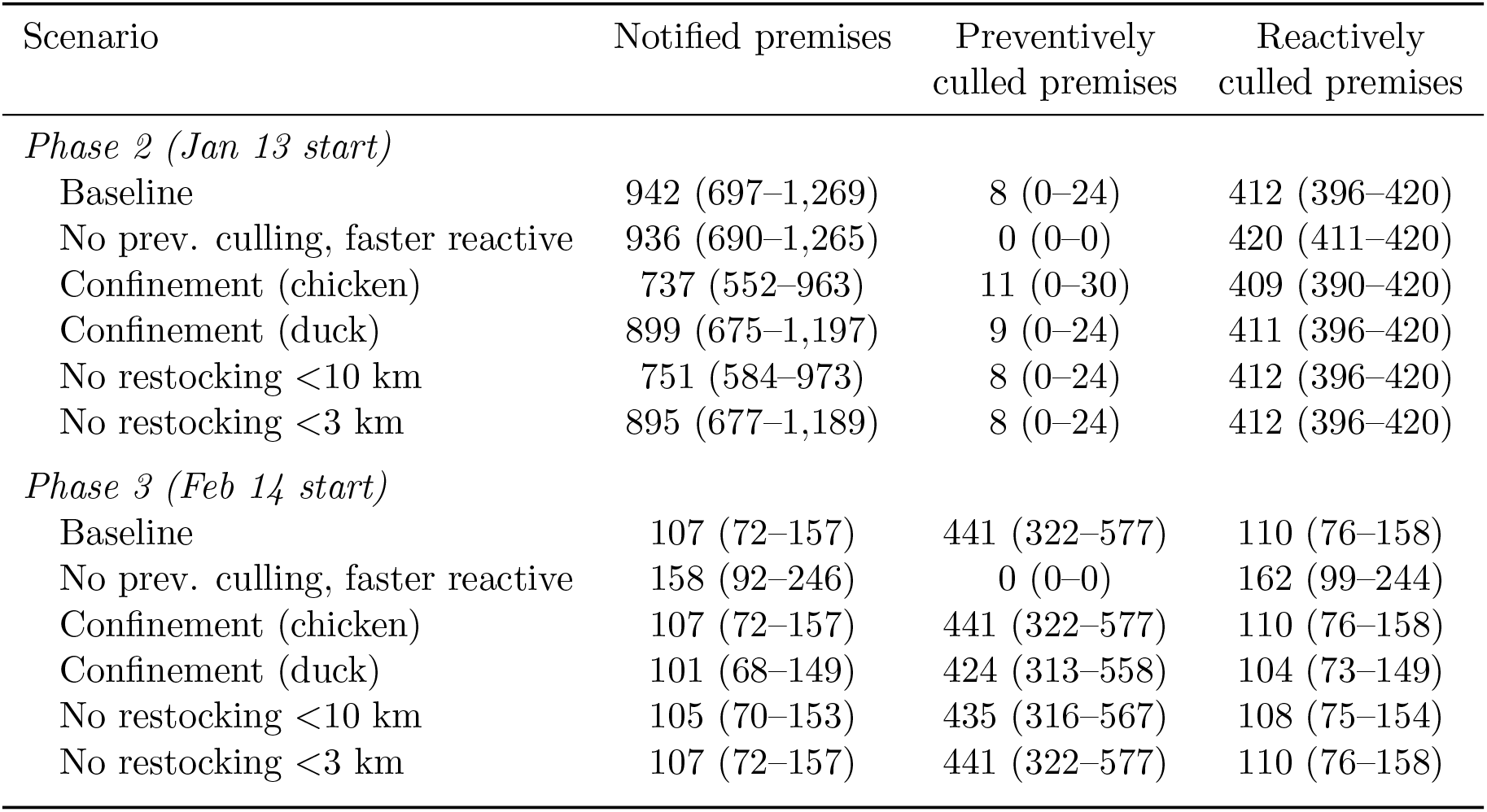
The median (95% PI) numbers of notified, preventively culled and reactively culled premises in our baseline and counterfactual model simulations over each 28-day projection window. Unlike the results in Fig. 2, these summary statistics reflect the total numbers of premises as opposed to the percentage change relative to the baseline scenario.

Since the PI spanned zero and the magnitude of the median effect was small, this result can be interpreted as representing no discernible difference between the projected effects of the two culling strategies over that 28-day window.

In contrast, introducing confinement on broiler stage two chicken premises was projected to lead to a large reduction in the number of notifications in the same period (with a median change of *−*21.6%, 95% PI (*−*38.2%*, −*7.6%)). Confinement on organic duck premises was projected to be less effective (*−*4.5%, 95% PI (*−*9.9%*, −*1.2%)) than confinement of broiler stage two chickens.

Restocking bans had large projected effects, with a median change in the projected number of notifications of *−*20.2% (95% PI (*−*25.4%*, −*14.5%)) when restocking was banned within ten km of notified premises and *−*5.0% (95% PI (*−*7.8%*, −*2.5%)) when restocking was banned within three km of notified premises (Fig. 2A).

In our simulations at the start of Phase 3, one month later, analogous analyses led to very different results (Fig. 2B). In particular, replacing preventive culling with faster reactive culling was projected to lead to a large increase in the number of notified premises (compared to no change in Phase 2). Specifically, as shown in Fig. 2B, the median change in the projected number of notifications was 45.0% (95% PI (17.1%, 84.2%)) in Phase 3 when preventive culling was replaced with faster reactive culling, suggesting that preventive culling would be very important to mitigate transmission in Phase 3. This demonstrates that the same control measures can have varying effects in different outbreak stages.

To understand why replacing preventive culling with faster reactive culling was projected to have little effect on the number of notified premises in Phase 2 but to increase it substantially in Phase 3, we compared the projections in these outbreak phases in more detail (Fig. 3, Table 2). In the Phase 2 window, the assumed culling capacity was *C*(*t*) = 15 premises per day, while the number of notified premises was high. Since, when multiple premises were due to undergo culling, reactive culling was prioritised over preventive culling, almost all of the available culling capacity was utilised for reactive culling. In the Phase 3 window, however, the culling capacity was *C*(*t*) = 100 premises per day and there were fewer notified premises. For that reason, in the baseline simulations a substantial amount of preventive culling was undertaken, and the counterfactual simulations with no preventive culling then led to a substantial projected increase in the number of notified premises. In other words, stopping preventive culling in Phase 3 was projected to allow substantial additional transmission (Fig. 2B), while in Phase 2 stopping the small amount of preventive culling that was being done had little effect (Fig. 2A).

**Figure 3:**
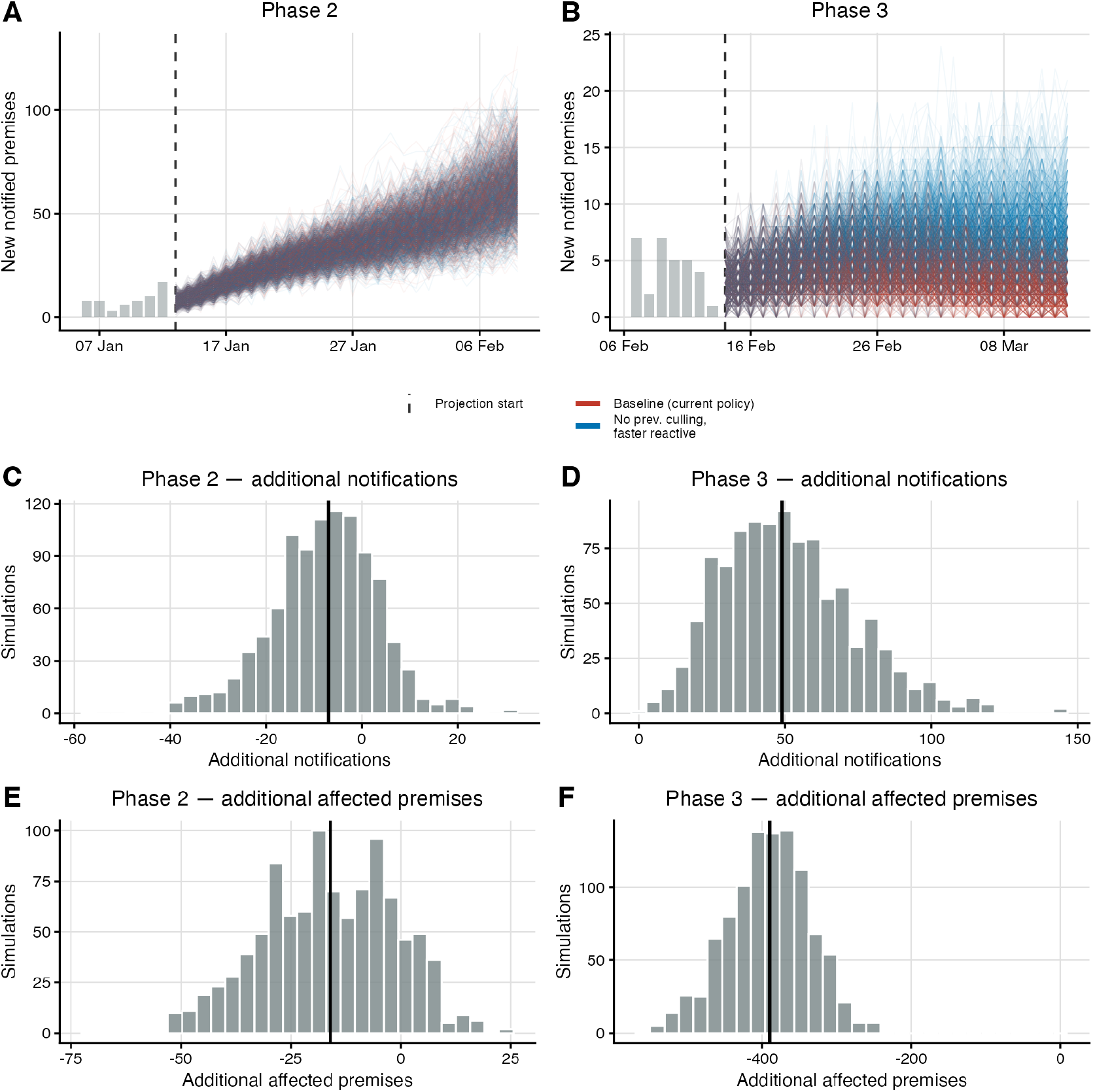
Comparison of projections under the baseline control policy against counterfactual scenarios with no preventive culling but faster reactive culling. (A) Individual simulated trajectories of the daily number of newly notified premises in the Phase 2 simulation window, under the baseline policy (red) and under the alternative strategy with no preventive culling but with faster reactive culling (blue). (B) Individual simulated trajectories of the daily number of newly notified premises in the Phase 3 window, under the baseline policy (red) and under the alternative strategy with no preventive culling but with faster reactive culling (blue). **(C)** Distribution of the difference in the number of notified premises between simulations under the alternative strategy relative to the baseline policy (alternative *−* baseline) in the Phase 2 window. **(D)** Distribution of the difference in the number of notified premises between simulations under the alternative strategy relative to the baseline policy (alternative *−* baseline) in the Phase 3 window. **(E)** Distribution of the difference in the number of affected premises between simulations under the alternative strategy relative to the baseline policy (alternative *−* baseline) in the Phase 2 window. **(F)** Distribution of the difference in the number of affected premises between simulations under the alternative strategy relative to the baseline policy (alternative *−* baseline) in the Phase 3 window. In panels A and B, each thin line represents one of 1,000 simulations per strategy, grey bars show the daily numbers of newly notified premises in the weeks immediately preceding the projection start and the dashed vertical line marks the projection start date. In panels C–F, negative values indicate that the alternative strategy reduces the relevant quantity, and the solid black line marks the median. Simulations were run for 28 days.

As described in Section 2.3, in addition to generating baseline model simulations (reflecting the control policy communicated to modelling teams at the decision points) and counterfactual scenarios, we also validated our model by running simulations reflecting the interventions that were ultimately implemented in the different outbreak phases. When we did this, we found that the model was able to recapitulate the outbreak dynamics that were actually observed in practice. Specifically, the observed number of notified premises lay within the 95% PI of the model simulations in both windows: there were actually 362 premises with notifications in the Phase 2 window (compared to a simulated median of 411, 95% PI (322, 528)) and 79 premises with notifications in the Phase 3 window (simulated median 108, 95% PI (72, 157)) (Fig. S2). In contrast, in our baseline model simulations, there were substantially more notified premises in the Phase 2 simulation window (median 942, 95% PI (697, 1,269); Table 2) than actually occurred, because the projections at decision time one assumed a lower culling capacity and no confinement.

### 3.2 The effectiveness of interventions depends on the outcome measure

In addition to the effectiveness of different control measures varying between stages of an outbreak, effectiveness also depends on the measure used to evaluate their success. We repeated the analyses presented in Fig. 2A,B, but instead quantifying the change in the total number of affected premises during the simulation period (Fig. 2C,D). We defined an “affected premises” as a premises on which either infections were notified or preventive culling was undertaken in the simulation period. We found that some interventions had similar projected effects on the change in the total number of notified premises and the total number of affected premises. For example, confinement of chickens in Phase 2 was projected to lead to a change in the total number of notified premises of *−*21.6% (95% PI (*−*38.2%*, −*7.6%)) and a change in the total number of affected premises of *−*21.1% (95% PI (*−*37.6%*, −*7.4%)). However, strategies involving stopping preventive culling had the potential to lead to very different projected effects when the two different outcome measures were considered. For example, in Phase 3, stopping preventive culling in favour of faster reactive culling led to a 45.0% (95% PI (17.1%, 84.2%)) change in the number of notified premises and a *−*71.4% (95% PI (*−*78.4%*, −*63.1%)) change in the number of affected premises. This highlights that interventions can be interpreted as beneficial or detrimental depending on whether the policy objective is to prioritise reducing transmission or to prioritise reducing the total number of premises that are either notified or preventively culled.

We note that, similarly to the results presented in Section 3.1, when we considered the total number of affected premises as the outcome metric of interest, we again found that the projected effectiveness of specific control measures depended on the outbreak stage. For example, unlike in Phase 3, in Phase 2 only a small decrease in the number of affected premises was projected if preventive culling was stopped in favour of faster reactive culling. As explained above in the context of the number of notified premises, this is because the large number of notified premises in the Phase 2 projections, in combination with the limited culling capacity, led to preventive culling only being undertaken on a small number of premises.

### 3.3 The relative priority of reducing transmission or limiting restrictions changes the preferred interventions

In Section 3.2, we compared the projected impacts of different control strategies in terms of the change in the number of notified premises and the change in the number of affected premises. If policy-makers choose which intervention to implement based on the first of these metrics, then they would effectively prioritise reducing transmission and assign no “cost” to control measures being in place. In contrast, choosing which intervention to implement according to the number of affected premises assigns the same cost to a notified premises and to a premises on which preventive culling is undertaken. In reality, policy-makers might weight the burden of interventions differently to the burden of infections. We therefore considered how the preferred control strategy changes as the cost of an intervention relative to a notified premises is varied.

In Phase 2, confinement of broiler stage two chickens was projected to be the most effective of the interventions considered (Fig. 2A,C). We therefore examined how the total outbreak cost, defined as the number of notified premises plus the weighted number of premises subject to confinement, varied with the assumed per-premises cost of confinement (Fig. 4A). As this cost increased, the projected benefit of confinement decreased. Confinement of chickens was projected to reduce the total outbreak cost relative to the baseline policy only if confining one premises was assumed to cost less than 0.095 times as much as one notified premises; for confinement of organic ducks, the corresponding threshold was 0.042. Above these thresholds, confinement was projected to increase the total outbreak cost. The lower threshold for ducks reflects the smaller number of notifications averted per confined premises.

**Figure 4:**
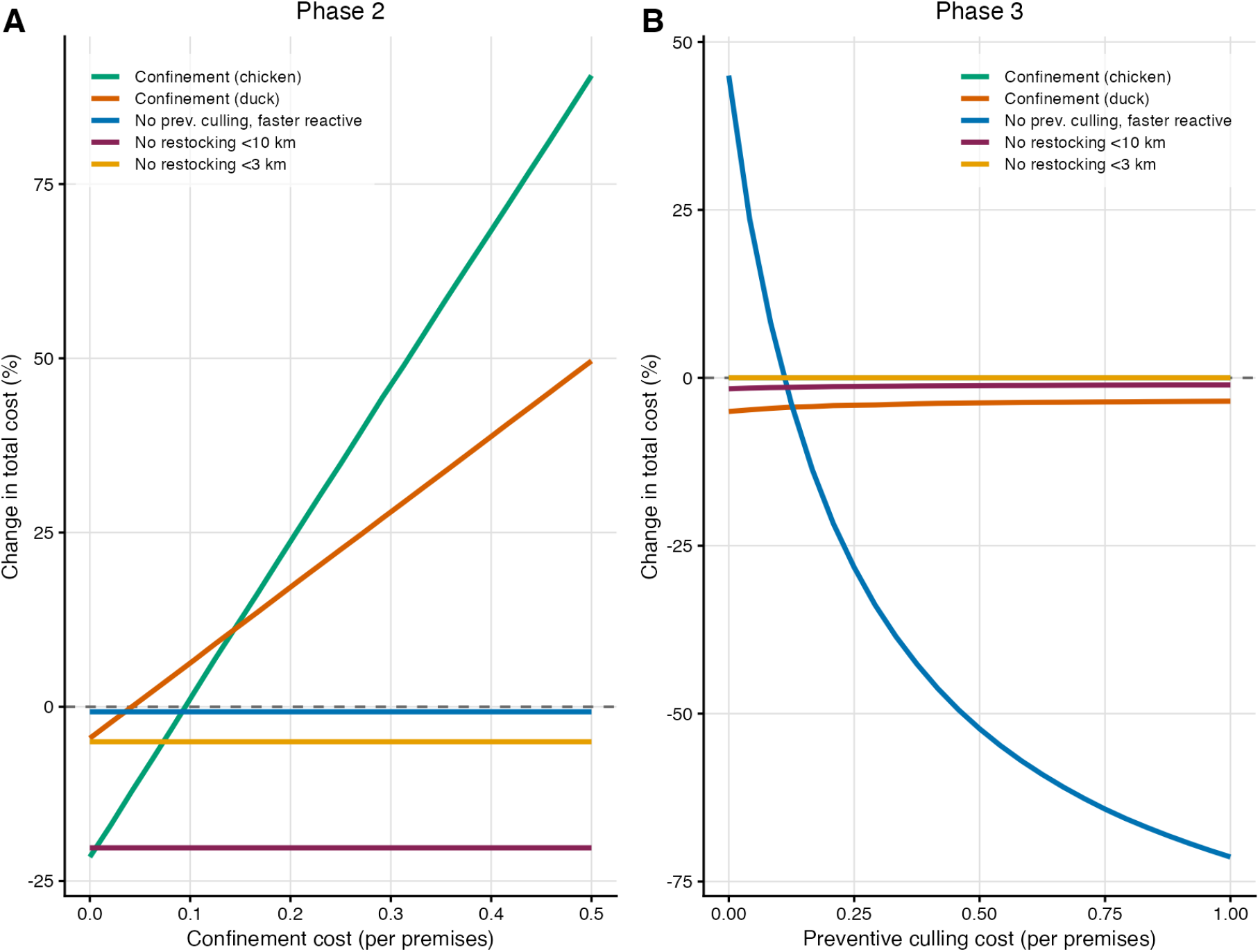
Projected change in the total outbreak cost under different control interventions as the relative cost of an intervention is varied. (A) Median percentage change in the total outbreak cost (relative to the baseline policy) in the Phase 2 window with different control interventions as the assumed per-premises confinement cost is varied. The total outbreak cost is defined as the number of notified premises plus the product of the number of premises subject to confinement and the per-premises cost of confinement. (B) Median percentage change in the total outbreak cost (relative to the baseline policy) in the Phase 3 window with different control interventions as the assumed per-premises preventive culling cost is varied. The total outbreak cost is defined as the number of notified premises plus the product of the number of preventively culled premises and the per-premises cost of preventive culling. In both panels, per-premises intervention costs are relative to the cost of one notified premises, the dashed horizontal line indicates no change relative to the baseline policy and simulations were run for 28 days.

In Phase 3, stopping preventive culling in favour of faster reactive culling was projected to increase the number of notified premises substantially (Fig. 2B) but to reduce the number of affected premises substantially (Fig. 2D). Whether this strategy is preferable therefore depends on the cost assigned to preventive culling on a premises relative to a notified premises (Fig. 4B). With the total outbreak cost defined as the number of notified premises plus the weighted number of premises on which preventive culling occurred, the median outbreak cost was lower under this strategy than under the baseline policy whenever a preventive cull was assumed to cost more than 0.11 times as much as a notified premises. Equivalently, retaining preventive culling was preferred only if policy-makers were willing to undertake around nine preventive culls to avert a single premises notification. Although these weights are not intended as an economic valuation, these results illustrate that the preferred strategy depends on the relative priorities assigned by policy-makers to reducing infections or limiting the extent of interventions.

## 4 Discussion

HPAI poses a substantial threat to the poultry industry worldwide. Mathematical models can be used to guide control interventions by enabling outbreak projections to be generated with a range of different control measures in place. For this purpose, we developed a spatially explicit, stochastic model of HPAI transmission with infections on poultry premises and used it to evaluate control strategies at two decision times during a synthetic HPAI outbreak. After fitting the model to the outbreak data using MCMC, we ran simulations over 28-day windows starting at the beginning of Phases 2 and 3 of the outbreak under alternative combinations of reactive culling, preventive culling, confinement and restocking bans. Use of the Sellke stochastic simulation algorithm ensured that the differences between outcomes of paired simulations reflected differences in the interventions that were implemented. We compared control strategies using different outbreak metrics: the number of notified premises, the number of affected premises and a weighted outbreak cost function under which the cost of notified infections was weighted differently to the cost of interventions.

Our first main finding is that the relative performance of different interventions changed during the outbreak. In particular, the intervention corresponding to the lowest value of the outbreak metric under consideration depended on the outbreak stage. A key factor driving this result is the difference in operational restrictions on interventions that were in place between outbreak phases. For example, at the first decision time, when there were large numbers of notified premises but a low culling capacity, replacing preventive culling with faster reactive culling led to essentially no change in the projected number of notifications in the simulation period (Figs 2A and 3C). One month later, when the culling capacity was higher and the number of notified premises had decreased, replacing preventive culling with faster reactive culling was projected to increase the number of notifications substantially (Figs 2B and 3D).

Our second main finding is that whether a specific intervention is successful or detrimental can depend on the metric used to evaluate its effectiveness. For example, preventive culling reduced the projected number of notifications in Phase 3 (Figs 2B and 3D), but did so by preemptively depopulating many premises. Consequently, preventive culling may be a beneficial control strategy if the policy objective is to reduce the number of notifications, but may not be beneficial if the policy objective is instead to reduce the number of affected premises (Figs 2D and 3F). We explored the trade-off between reducing transmission and maintaining low intervention costs in Fig. 4. We found, for example, that the culling strategy corresponding to the lower cost depended on whether or not the per-premises cost of preventive culling exceeded a specific threshold (Fig. 4B).

Our first finding highlights the benefits of so-called “adaptive management” strategies under which control interventions are updated during an outbreak according to the epidemiological situation and associated uncertainty (Merl et al., 2009; Shea et al., 2014; Probert et al., 2018; Atkins et al., 2020). Our second finding highlights the importance of policy-makers stating their objectives explicitly (Probert et al., 2016). If this is done, epidemiological models can in principle be used to identify optimal intervention policies. We note that, in scenarios in which policy-makers are not explicit about the policy objectives, models can be used to solve the reverse problem (i.e., to determine the importance that policy-makers have implicitly assigned to preventing infections as opposed to reducing disease control costs). We also note that policymakers’ objectives may shift during an outbreak: for example, in the earliest stages of an emerging outbreak, policy-makers might seek to eliminate the pathogen, whereas later when there are large numbers of cases they might instead seek to maintain case numbers below a specified (non-zero) threshold (Thompson et al., 2026). Quantitative analyses that seek to identify optimal control strategies must be adapted accordingly.

Our findings are relevant to policy decisions about HPAI control that are currently being made in countries worldwide. Preventive culling can reduce transmission, but its desirability depends on both operational feasibility and the extent to which onwards transmission occurs from premises with infections (as opposed to, for example, spillovers from a wild bird reservoir driven by environmental factors that are independent of existing infections on poultry premises (Hayes et al., 2026b)). In France, a reduction in duck farm density (for example, by preventive culling) has been proposed to reduce transmission (Vergne et al., 2024). In the same country, mandatory vaccination of commercial ducks has been introduced (Guinat et al., 2025), and vaccination is also being considered elsewhere (Department for Environment, Food and Rural Affairs, 2026). Although vaccination was not evaluated in the present analysis, in principle vaccination could be incorporated as an additional intervention in our transmission model. We also note that our findings extend beyond HPAI to a wide range of agricultural pathogens. For example, spatially explicit premises-level models have been used to evaluate control measures against diseases such as foot-and-mouth disease (Keeling et al., 2001; Tildesley et al., 2006, 2009) and African swine fever (Hayes et al., 2021; Dankwa et al., 2022). For a wide range of pathogens, both the outbreak stage (accounting for different operational constraints on interventions) and the policy objective are likely to impact the effectiveness of control measures.

As with any modelling analysis, our study has limitations. Perhaps the most substantial limitation is that this research was based on analysing a hypothetical HPAI outbreak, rather than a real-world outbreak. As such, quantitative values in this work should not be transferred directly to inform real-world control policy. We also fitted the model to data from the entire outbreak before using model simulations to generate projections in different outbreak phases. Fitting to data from the entire outbreak provided a controlled retrospective comparison in which the model was not sequentially updated between decision times, enabling the simulations to reflect differences between the interventions and assumed culling capacity being considered at the decision times rather than differences in model structure or parameterisation. Our analysis should therefore not be interpreted as a reconstruction of the policy-relevant analyses that would have been generated in real-time. Finally, in our weighted-objective analysis presented in Fig. 4, we considered a simple linear cost function. Use of our model to guide policy in a real-world outbreak requires a more detailed epidemiological-economic analysis to be undertaken, to ensure that decisions are based on realistic estimates of the value of reducing transmission and the costs of interventions.

Despite these limitations, we were able to demonstrate two key disease control principles that should be considered further and tested more widely. We showed that intervention evaluation during an HPAI outbreak is a sequential decision problem rather than a one-off optimisation: the preferred intervention strategy can change as an outbreak progresses. We also demonstrated the importance of policy objectives being chosen carefully and stated explicitly. Flexible responses to agricultural outbreaks that are evaluated against clearly stated policy objectives, and reassessed as the epidemiological situation and operational constraints change, have the potential to limit the detrimental economic and animal health impacts of outbreaks.

## Acknowledgements

We thank the organisers of the HPAI Modelling Challenge (WiLiMan-ID consortium) for designing the challenge scenario and providing the epidemiological data underlying our analyses. Thanks to the Infectious Disease Modelling group in the Mathematical Institute at the University of Oxford for helpful discussions throughout the project. We also acknowledge the use of the University of Oxford’s Advanced Research Computing (ARC) facility (https://doi.org/10.5281/zenodo.22558). F.B., E.M.H., M.J.T. and R.N.T. are supported by the Biotechnology and Biological Sciences Research Council (BBSRC) grant *Modelling to inform interventions during Highly Pathogenic Avian Influenza outbreaks in Great Britain* (grant number BB/X016137/1).

E.M.H. is affiliated to the NIHR Health Protection Research Unit in Emerging and Zoonotic Infections (NIHR HPRU-EZI) (NIHR207393) at the University of Liverpool in partnership with the UK Health Security Agency (UKHSA), in collaboration with Liverpool School of Tropical Medicine, London School of Hygiene and Tropical Medicine and The University of Oxford.

E.M.H. is funded by The Pandemic Institute, formed of seven founding partners: The University of Liverpool, Liverpool School of Tropical Medicine, Liverpool John Moores University, Liverpool City Council, Liverpool City Region Combined Authority, Liverpool University Hospital Foundation Trust, and Knowledge Quarter Liverpool (E.M.H. is based at The University of Liverpool). The views expressed are those of the author(s) and not necessarily those of the NIHR, the Department of Health and Social Care, UKHSA or The Pandemic Institute.

## Data and code availability

Computing code for running the model and estimating its parameter values from outbreak data is available at https://github.com/fannybergstrom/hpai_challenge. The epidemiological data were provided by the organisers of the HPAI Modelling Challenge and are included in this repository.

## Authors’ contributions

Fanny Bergström: Conceptualization, Formal analysis, Software, Project administration, Writing – original draft, Writing – review & editing. Edward M. Hill: Conceptualization, Funding acquisition, Writing – review & editing. Cecilia Rago: Visualization, Writing – review & editing. Christopher N. Davis: Conceptualization, Writing – review & editing. Michael J. Tildesley: Conceptualization, Funding acquisition, Writing – review & editing. Robin N. Thompson: Conceptualization, Funding acquisition, Writing – original draft, Writing – review & editing.

## Declaration of competing interest

The authors declare that they have no known competing financial interests or personal relationships that could have appeared to influence the work reported in this paper.

## Appendix

### A Model parameters

Here, we summarise the parameters of our transmission model. In Table S1, we provide parameter definitions, units and whether the parameter was inferred from the outbreak data (see Section 2.2) or fixed at values from the literature (with references provided for those parameters). Throughout, the index *k* denotes the production type of the susceptible (infectee) premises *i* and *l* denotes the production type of the infectious or notified (infector) premises *j*. The priors used for the inferred parameters are given in Table S2, and posterior–prior comparisons are shown in Fig. S1.

**Table S1:**
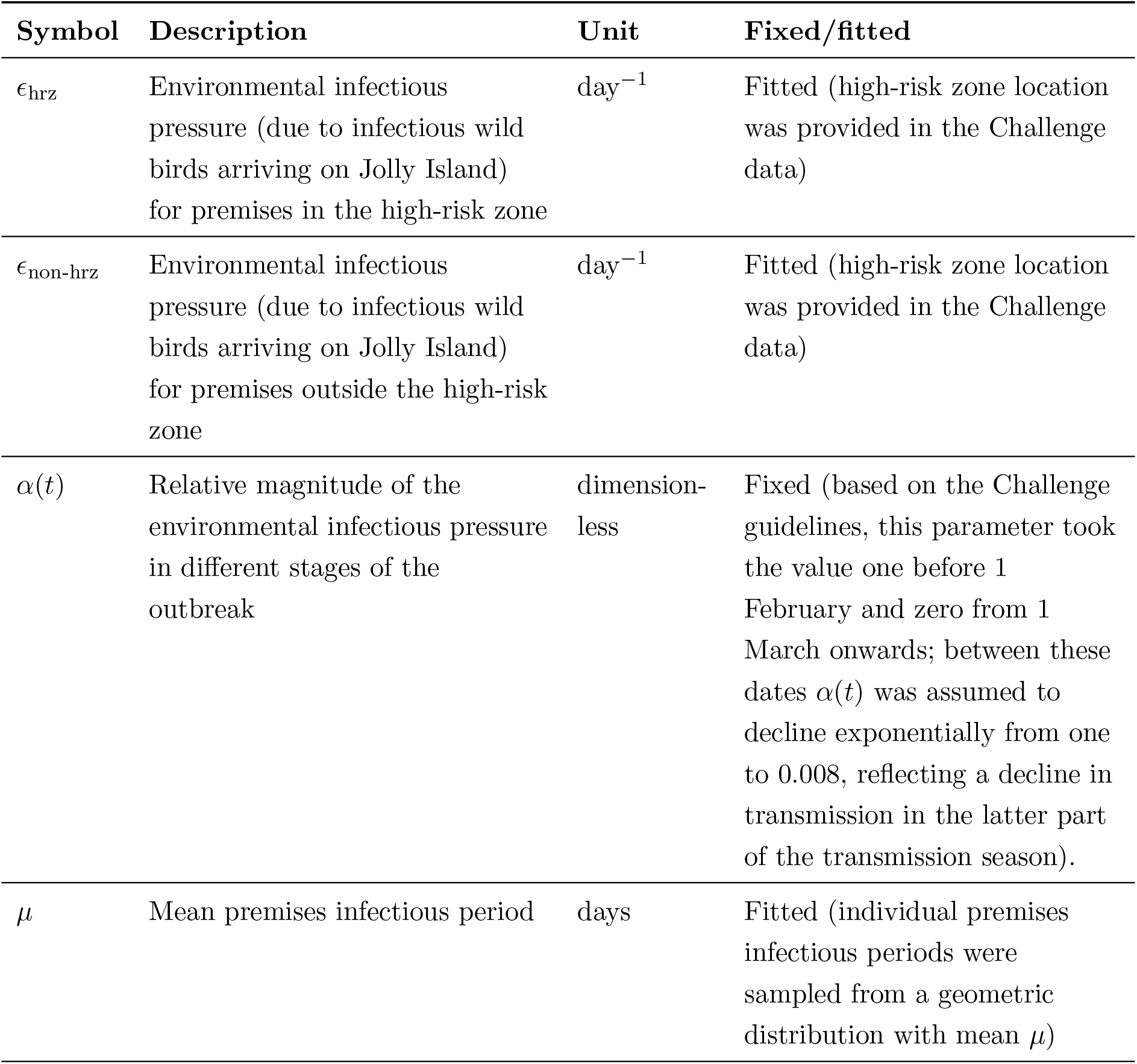
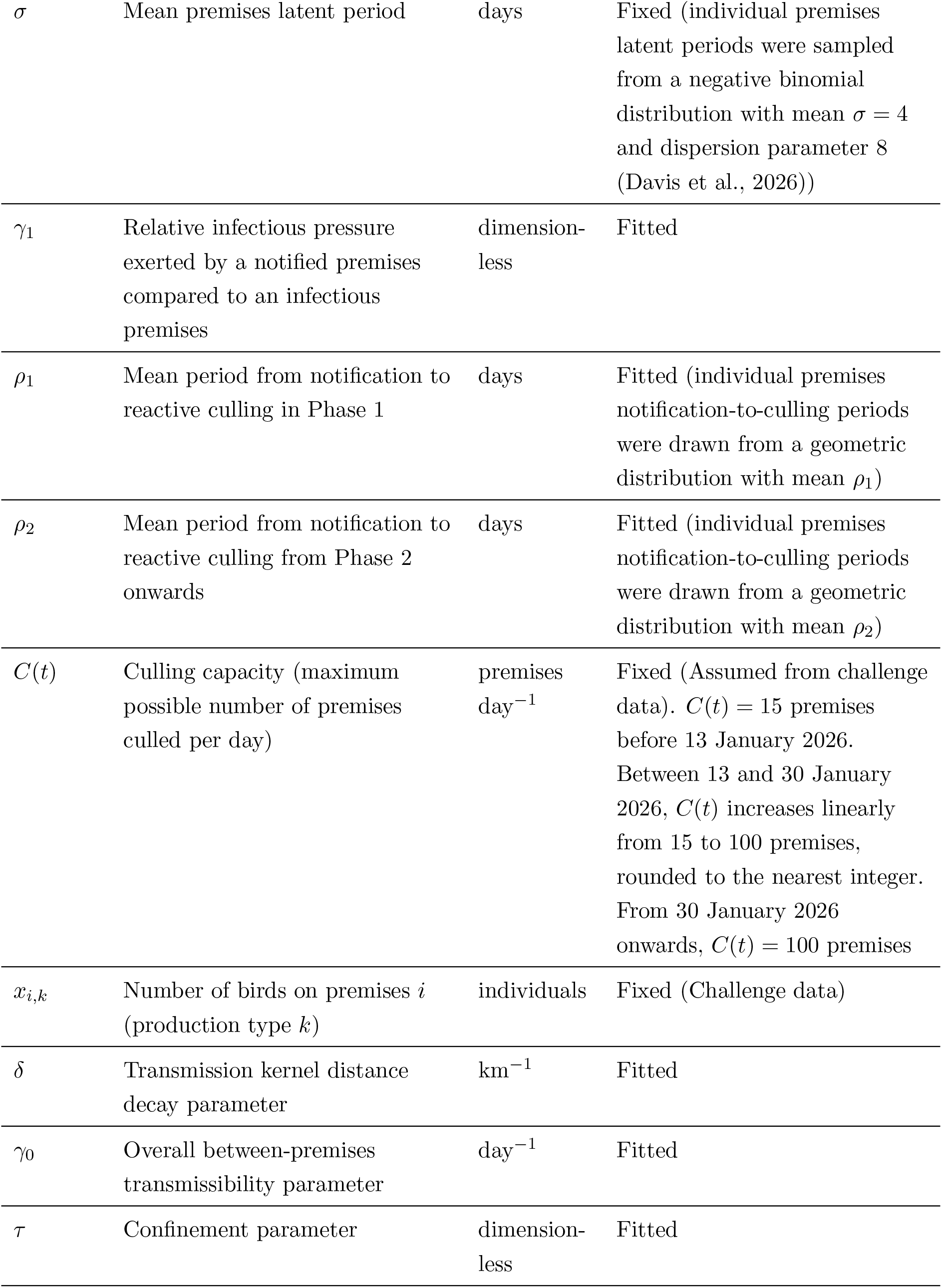
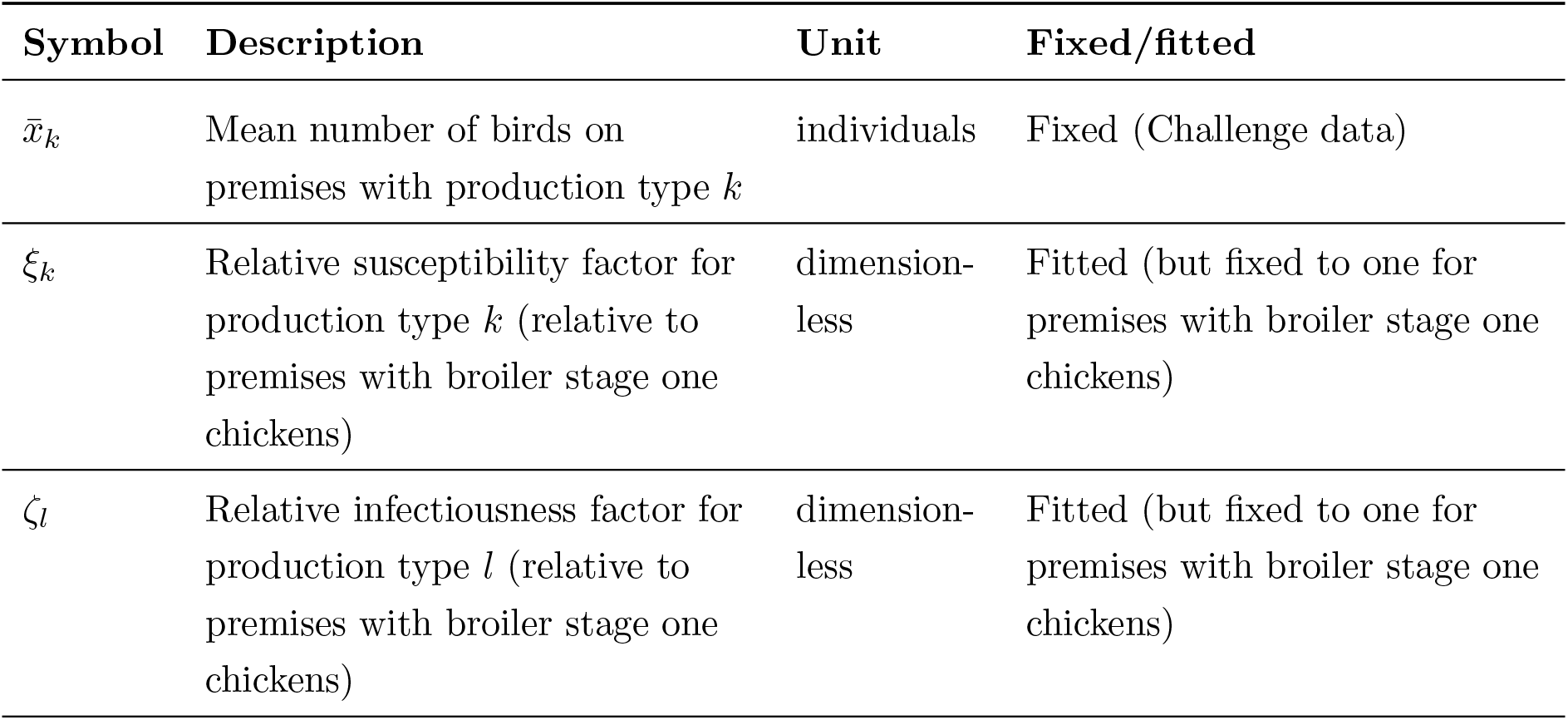
Model parameters, their definitions, units and whether the parameter was inferred from the outbreak data or fixed. Literature sources are provided for the fixed values that were not provided by the Challenge organisers.

| Symbol | Description | Unit | Fixed/fitted |
| --- | --- | --- | --- |
| $\epsilon_{\text{hrz}}$ | Environmental infectious pressure (due to infectious wild birds arriving on Jolly Island) for premises in the high-risk zone | $\text{day}^{-1}$ | Fitted (high-risk zone location was provided in the Challenge data) |
| $\epsilon_{\text{non-hrz}}$ | Environmental infectious pressure (due to infectious wild birds arriving on Jolly Island) for premises outside the high-risk zone | $\text{day}^{-1}$ | Fitted (high-risk zone location was provided in the Challenge data) |
| $\alpha(t)$ | Relative magnitude of the environmental infectious pressure in different stages of the outbreak | dimensionless | Fixed (based on the Challenge guidelines, this parameter took the value one before 1 February and zero from 1 March onwards; between these dates $\alpha(t)$ was assumed to decline exponentially from one to 0.008, reflecting a decline in transmission in the latter part of the transmission season). |
| $\mu$ | Mean premises infectious period | days | Fitted (individual premises infectious periods were sampled from a geometric distribution with mean $\mu$) |
| $\sigma$ | Mean premises latent period | days | Fixed (individual premises latent periods were sampled from a negative binomial distribution with mean $\sigma = 4$ and dispersion parameter 8 (Davis et al., 2026)) |
| $\gamma_1$ | Relative infectious pressure exerted by a notified premises compared to an infectious premises | dimensionless | Fitted |
| $\rho_1$ | Mean period from notification to reactive culling in Phase 1 | days | Fitted (individual premises notification-to-culling periods were drawn from a geometric distribution with mean $\rho_1$) |
| $\rho_2$ | Mean period from notification to reactive culling from Phase 2 onwards | days | Fitted (individual premises notification-to-culling periods were drawn from a geometric distribution with mean $\rho_2$) |
| $C(t)$ | Culling capacity (maximum possible number of premises culled per day) | premises day<sup>-1</sup> | Fixed (Assumed from challenge data). $C(t) = 15$ premises before 13 January 2026. Between 13 and 30 January 2026, $C(t)$ increases linearly from 15 to 100 premises, rounded to the nearest integer. From 30 January 2026 onwards, $C(t) = 100$ premises |
| $x_{i,k}$ | Number of birds on premises $i$ (production type $k$) | individuals | Fixed (Challenge data) |
| $\delta$ | Transmission kernel distance decay parameter | km<sup>-1</sup> | Fitted |
| $\gamma_0$ | Overall between-premises transmissibility parameter | day<sup>-1</sup> | Fitted |
| $\tau$ | Confinement parameter | dimensionless | Fitted |
| $\bar{x}_k$ | Mean number of birds on premises with production type $k$ | individuals | Fixed (Challenge data) |
| $\xi_k$ | Relative susceptibility factor for production type $k$ (relative to premises with broiler stage one chickens) | dimensionless | Fitted (but fixed to one for premises with broiler stage one chickens) |
| $\zeta_l$ | Relative infectiousness factor for production type $l$ (relative to premises with broiler stage one chickens) | dimensionless | Fitted (but fixed to one for premises with broiler stage one chickens) |

**Table S2:** Priors used for the fitted parameters of the transmission model. We list the relevant poultry types for the *ξ_k_* and *ζ_l_* variables. Note that for the gamma distributed priors, we use the (shape, rate) parameterisation.

| Parameter | Prior distribution |
| --- | --- |
| $\gamma_0$ (overall transmissibility) | Gamma(3, 136) |
| $\gamma_1$ (relative infectious pressure of $N$ vs $I$) | Beta(7, 4) |
| $\epsilon_{\text{hrz}}$ (environmental infectious pressure in HRZ) | Lognormal(log 0.0009, 0.6) |
| $\epsilon_{\text{non-hrz}}$ (environmental infectious pressure outside HRZ) | Lognormal(log 0.00018, 0.6) |
| $\tau$ (confinement parameter) | Beta(8, 2) |
| $\delta$ (transmission kernel parameter) | Lognormal(log 0.17, 0.30) |
| $\mu$ (mean infectious period) | Gamma(16, 4) |
| $\rho_1$ (mean reactive culling delay in Phase 1) | Gamma(17, 5.9) |
| $\rho_2$ (mean reactive culling delay from Phase 2 onwards) | Gamma(36, 15) |
| $\xi_2$ (susceptibility, broiler stage two chickens) | Lognormal(log 1.5, 0.65) |
| $\xi_3$ (susceptibility, layer chickens) | Lognormal(log 0.78, 0.65) |
| $\xi_4$ (susceptibility, conventionally reared ducks) | Lognormal(log 0.13, 0.60) |
| $\xi_5$ (susceptibility, organic ducks) | Lognormal(log 0.40, 0.65) |
| $\zeta_2$ (infectiousness, broiler stage two chickens) | Lognormal(log 0.70, 0.65) |
| $\zeta_3$ (infectiousness, layer chickens) | Lognormal(log 0.38, 0.70) |
| $\zeta_4$ (infectiousness, conventionally reared ducks) | Lognormal(log 0.32, 0.70) |
| $\zeta_5$ (infectiousness, organic ducks) | Lognormal(log 0.40, 0.70) |

### B Parameter estimates and convergence diagnostics

Here, we report the posterior estimates and MCMC convergence diagnostics for the fitted model parameters. In all our model simulations, we sampled the parameters used in each simulation from the joint posterior distribution. Marginal posterior medians and 95% CrIs for all estimated parameters are stated in Table S3. We show the marginal posteriors for each fitted parameter alongside the priors in Fig. S1. We estimated the posterior densities using Gaussian kernel density estimation (KDE), computed with stats::density() in R (called via ggplot2::geom density()) using its default settings: a Gaussian kernel and a bandwidth using the bw.nrd0 rule-of-thumb selector (Silverman, 1986).

**Figure S1:**
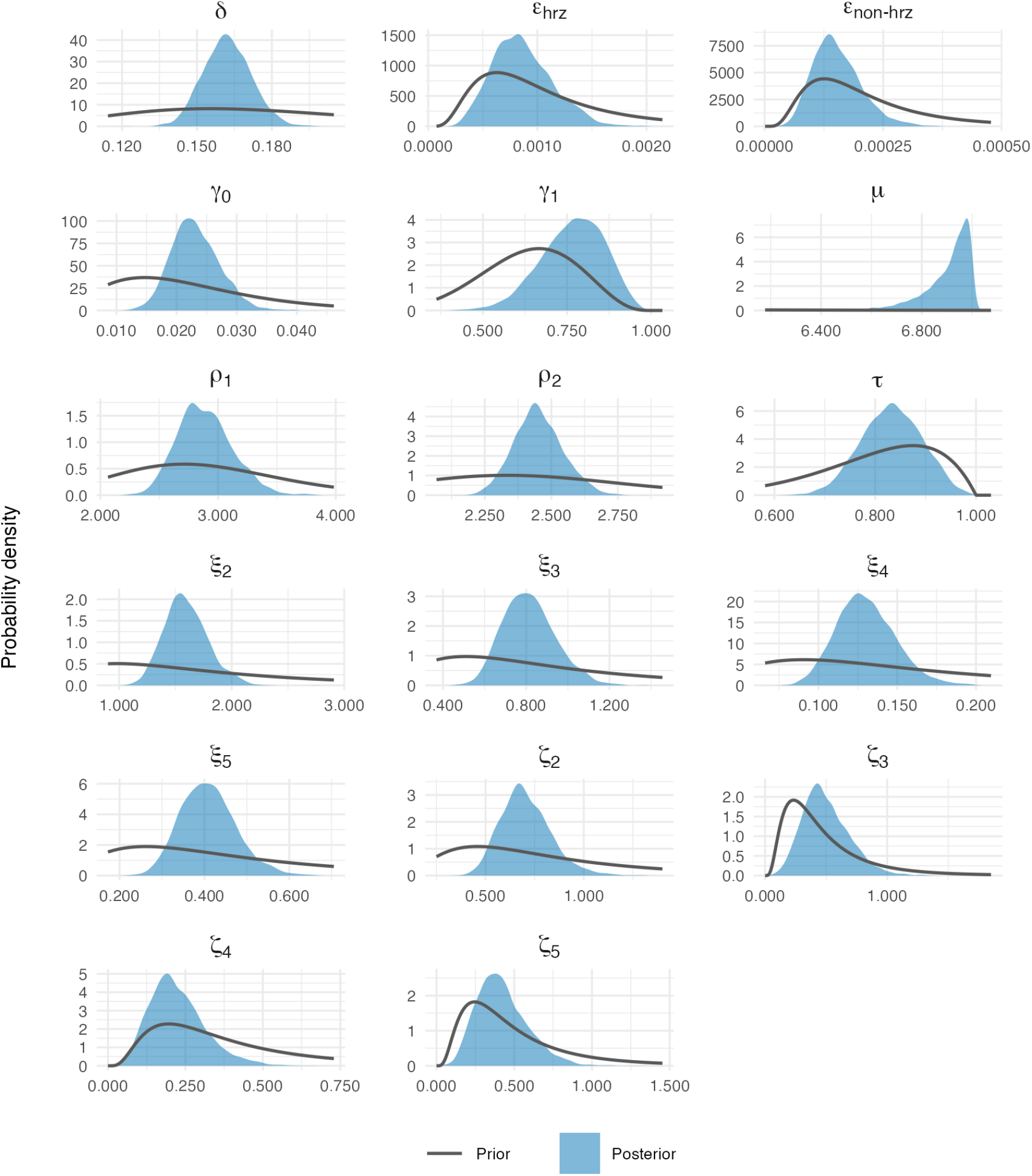
Posterior (blue) and prior (grey) distributions for the fitted model parameters. The plotted posterior distributions are kernel density estimates over the full set of posterior draws. The priors are the probability distributions specified in Table S2. We set the x-axes ranges to enable the posterior distributions to be visualised clearly.

**Table S3:** Posterior medians (and 95% credible intervals) and MCMC diagnostics for the fitted model parameters. Gelman-Rubin statistic (*R̂*) and effective sample sizes are calculated as described by Vehtari et al. (2021). Values of *R̂* close to one indicate adequate mixing and convergence across different chains, with *R̂≤* 1.01 taken as evidence of satisfactory convergence. The effective sample size (ESS) measures the amount of effectively independent information contained in the autocorrelated MCMC draws; values of ESS *≥* 400 are taken to indicate adequate sampling efficiency and precision. All inferred parameters meet these thresholds.

| Parameter | Median (95% CrI) | $\hat{R}$ | ESS (bulk) | ESS (tail) |
| --- | --- | --- | --- | --- |
| $\gamma_0$ | 0.023 (0.016, 0.032) | 1.003 | 3405 | 2997 |
| $\gamma_1$ | 0.774 (0.567, 0.924) | 1.001 | 6098 | 2224 |
| $\epsilon_{\text{hrz}}$ | 0.0008 (0.0004, 0.0014) | 1.000 | 5386 | 2844 |
| $\epsilon_{\text{non-hrz}}$ | 0.00015 (0.00007, 0.00027) | 1.001 | 6058 | 2816 |
| $\tau$ | 0.833 (0.716, 0.944) | 1.000 | 4228 | 2436 |
| $\delta$ | 0.162 (0.145, 0.181) | 1.002 | 5069 | 2548 |
| $\mu$ | 6.934 (6.649, 6.998) | 1.001 | 4817 | 1606 |
| $\rho_1$ | 2.873 (2.479, 3.389) | 1.001 | 8042 | 3094 |
| $\rho_2$ | 2.447 (2.281, 2.641) | 1.001 | 8007 | 2756 |
| $\xi_2$ (broiler stage two chickens) | 1.579 (1.253, 2.020) | 1.000 | 3604 | 3398 |
| $\xi_3$ (layer chickens) | 0.807 (0.595, 1.086) | 1.000 | 4683 | 3217 |
| $\xi_4$ (conventionally reared ducks) | 0.129 (0.097, 0.171) | 1.001 | 4170 | 2827 |
| $\xi_5$ (organic ducks) | 0.406 (0.295, 0.552) | 1.001 | 4513 | 3011 |
| $\zeta_2$ (broiler stage two chickens) | 0.693 (0.490, 0.995) | 1.001 | 4541 | 3390 |
| $\zeta_3$ (layer chickens) | 0.472 (0.189, 0.932) | 1.002 | 5060 | 2123 |
| $\zeta_4$ (conventionally reared ducks) | 0.211 (0.085, 0.434) | 1.003 | 5490 | 2140 |
| $\zeta_5$ (organic ducks) | 0.402 (0.166, 0.791) | 1.000 | 5494 | 2697 |

### C Model validation against outbreak data

To assess whether the fitted model could reproduce the observed outbreak dynamics, we compared the daily number of notified premises in model simulations against those observed in the dataset in each of the two 28-day simulation windows. When we ran these simulations, we implemented the interventions that were known to be in place during the simulation windows. Specifically, the culling capacity followed the realised increase from 15 to 100 premises per day between 13 and 30 January 2026 (Table S1) and confinement was implemented on broiler stage two chicken and organic duck premises according to the dates on which it was in place. In the Phase 2 window, these simulations therefore differ from our baseline modelling projections in which we fixed the culling capacity at *C*(*t*) = 15 premises per day and did not implement confinement (Section 2.3, Table 2). We ran 1,000 simulations per window. Each simulation used a parameter set that was sampled from the joint posterior distribution. For each window, we summarised the simulated total number of notified premises by its median and central 95% PI, and compared this with the total in the dataset.

The number of notified premises in the dataset lay within the 95% PI of the simulations in both windows. In the Phase 2 window, there were 362 premises with notified infections, compared to a median of 411 (95% PI (322,528)) in the simulations. The simulations reproduced the timing of the peak in daily notifications in late January 2026 and the subsequent decline.

The daily incidence of notifications was mostly contained within the 95% PI each day in the simulations. In the Phase 3 window, there were 79 notified premises in the dataset, compared to 108 (95% PI (72,157)) in the simulations; the median of three to four notified premises per day in the simulations exceeded the observed daily counts on most days, although the latter was again within the 95% PI each day.

**Figure S2:**
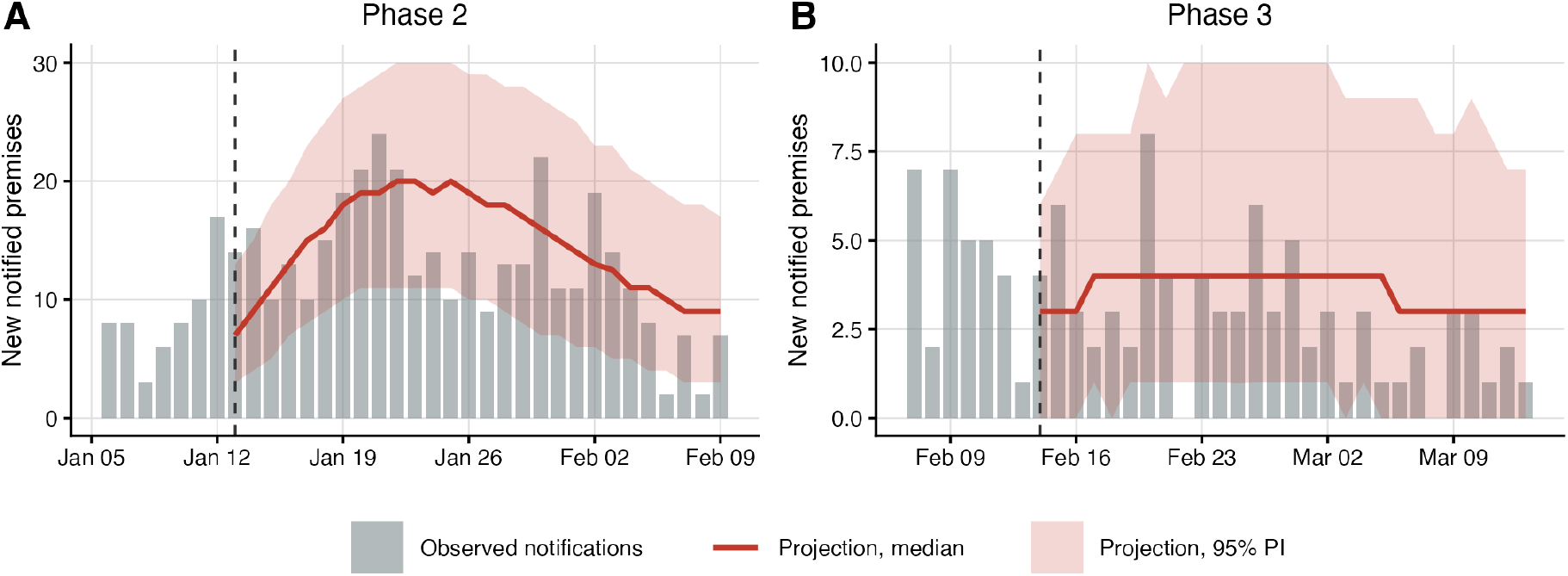
Verification that the transmission model can reproduce the observed outbreak data. Daily numbers of newly notified premises (grey bars) for the weeks immediately preceding each projection window and over the window itself, together with the median (red line) and pointwise central 95% PI (shaded) obtained from 1,000 model simulations in which the culling capacity and confinement were set according to the interventions that were in place in those windows. (A) Data and simulation results for the Phase 2 window, which began on 13 January 2026. (B) Data and simulation results for the Phase 3 window, which began on 14 February 2026. The dashed vertical line marks the start of each simulation window.

## Notes

### Competing Interest Statement

The authors have declared no competing interest.

